# A new *Serratia* taxon found in a trombidiform mite shows genomic evidence of host adaptation but is phylogenetically distinct from *Serratia symbiotica*

**DOI:** 10.64898/2026.11.03.764106

**Authors:** Jia Chang, Ary A. Hoffmann, Neil D. Young, Sunita B. Sumanam, Thomas L. Schmidt, Joshua A. Thia

## Abstract

The evolutionary transition from free-living bacteria to host-adapted symbionts is typically accompanied by substantial genomic changes. Within the genus *Serratia*, this transition has so far been documented only in the aphid-associated species, *Serratia symbiotica*, whereas all other *Serratia* species are free-living. In this study, we identify a putatively new *Serratia* taxon using metagenomic approaches in the trombidiform mite, *Halotydeus destructor*. We provide genomic evidence for a possible transition to a host-adapted lifestyle in this new *Serratia* sp., which sits phylogenetically within a clade of free-living *Serratia* species. The genomic signatures of evolving toward a host-adapted lifestyle include a reduced genome size and coding capacity, an elevated transposase fraction, and detectable chromosomal rearrangements, relative to other free-living *Serratia* species. However, the species does not show the same structure of genome reduction as *S. symbiotica*. Functional gene content analyses including COG categories and virulence factor repertoires also showed that our new *Serratia* sp. had a distinct gene profile from both free-living *Serratia* and *S. symbiotica*. Additional genomic screening suggests that infections of the new *Serratia* sp. are common in wild populations of *H. destructor* across its invasive Australian range, warranting future investigations on the possible role this symbiont plays in the evolutionary ecology of its mite host. Our work not only highlights a potential parallel transition to a host-adapted lifestyle in the *Serratia* genus, but also represents the first characterisation of a possible host-adapted *Serratia* taxon in a trombidiform mite.

## Introduction

Many arthropods carry bacterial symbionts that have undergone major adaptive changes to integrate into the biology of their hosts (McCutcheon and Moran 2007; Weiss et al. 2008; Powell et al. 2016). The most well-known examples include bacteria that have become involved in nutritional symbioses. For example, *Blochmannia* in carpenter ants provides essential amino acids to its host (Feldhaar et al. 2007). Similarly, *Buchnera* in aphids provision their hosts with essential amino acids and vitamins (Douglas 1998). Such symbionts represent evolutionary extremes in which bacterial genomes have become highly streamlined and reduced as a result of obligate symbiosis with their insect host. However, other symbiotic bacteria exist on a continuum from free-living to host-adapted. *Burkholderia* in bean bugs has demonstrated positive effects on pesticide resistance and metabolite recycling, but it retains its capacity to be free-living (Kikuchi et al. 2012; Ohbayashi et al. 2019). In scale insects, several bacterial species are believed to have recently transitioned from free-living to host-adapted lifestyles, including those in the genera *Serratia, Pantoea, Enterobacter*, and *Kosakonia* (Palanichamy et al. 2025). Experimental data remains the most important way of demonstrating critical associations between arthropods and their host-adapted bacterial symbionts (Feldhaar et al. 2007; Akman Gündüz and Douglas 2009). Nevertheless, genomic signatures of host adaptation provide a valuable tool for placing bacterial symbionts on the free-living versus host-adapted continuum in the absence of experimental evidence.

A classic signature in the transition to a host-adapted lifestyle is genome reduction, which includes decreasing genome size, gene loss and pseudogenization, and increased AT-bias (McCutcheon and Moran 2012). One reason for such reduction is genomes are selected for increased streamlining in host-adapted bacterial symbionts. As bacteria become increasingly integrated into the biology of their host, there is weakened selection to maintain genes that are needed for a free-living lifestyle, but ongoing selection on crucial genes and those important to their symbiosis (Toft and Andersson 2010; McCutcheon and Moran 2012). The genome reduction process is well exemplified by *Carsonella*, a psyllid symbiont that has undergone extensive gene loss, resulting in one of the smallest and most AT-rich bacterial genomes that nevertheless retains genes involved in essential amino acid biosynthesis (Sloan and Moran 2012).

The transition to host adaptation is also associated with increased activity of mobile elements (Moran and Plague 2004; Plague et al. 2008). One hypothesis for such proliferation is the reduced selection on CRISPR-Cas systems that keep mobile elements in check as bacteria become increasingly host-adapted (Siozios et al. 2024). The proliferation of mobile elements can promote gene inactivation and loss, facilitate genome rearrangements, and alter the expression of neighbouring genes, thereby facilitating genome restructuring and reduction (Moran and Plague 2004). For example, recently evolved endosymbiotic *Arsenophonus* lineages show extensive accumulation of mobile genetic elements (Siozios et al. 2024). Similarly, *Sodalis glossinidius*, a tsetse fly endosymbiont at an early stage of transition from a free-living to a host-dependent lifestyle, exhibits extensive proliferation of insertion sequences associated with early symbiosis (Belda et al. 2010).

Virulence factor genes provide another lens through which to assess transitions from free-living to host-adapted lifestyles. Overall, virulence-associated genes are lost or inactivated alongside other genes during genome reduction. However, some ancestral virulence factors are retained because they contribute to host colonization (Dale and Moran 2006; McCutcheon et al. 2019). In that case, virulence factor profiles might shift away from ancestral free-living composition to a new profile associated with host adaptation. Although few studies have documented such a shift during the transition from free-living to host-adapted lifestyles, changes in virulence factor repertoires associated with adaptation to different hosts have been reported in *Pseudomonas*, *Salmonella*, and *Hamiltonella* (Hulin et al. 2018; Den Bakker et al. 2011; Rollat-Farnier et al. 2015).

The genus *Serratia* comprises Gram-negative bacteria (Grimont et al. 1977) that have established different associations with arthropod hosts. Most *Serratia* species are free-living and can be found in diverse environments including water and soil, as well as inside animals and plants (Petersen and Tisa 2013). Some free-living *Serratia* species can be symbiotic and persist in the host as commensal symbionts or opportunistic pathogens. For example, *Serratia odorifera* is stably associated with the midgut of *Aedes aegypti* across multiple developmental stages (Apte-Deshpande et al. 2012), *Serratia nematodiphila* forms a stable association with entomopathogenic nematodes (Kwak et al. 2015), and *Serratia marcescens* strains can establish persistent colonization in multiple insect hosts including mosquitos, honey bees, and triatomine bugs (Chen et al. 2017; Raymann et al. 2018; Azambuja et al. 2004). Despite their symbiotic associations, these species retain large genomes typical of free-living *Serratia* and show no evidence of the genomic changes associated with host adaptation.

Only a single species of *Serratia*, *S. symbiotica*, is considered to be host-adapted. *Serratia symbiotica* is characterised by significant genome reduction and genomic rearrangement during specialisation to aphid hosts (Burke and Moran 2011; Manzano-Marín and Latorre 2016). *Serratia symbiotica* strains are particularly associated with aphids, but have evolved diverse associations ranging from gut pathogens to facultative mutualists and obligate mutualists (Perreau et al. 2021; Monnin et al. 2020). For example, a gut pathogenic strain (CWBI-2.3) in a relatively early stage of host adaptation can impose fitness costs in *Aphis fabae* under certain environmental conditions (Pons et al. 2019). In contrast, a facultative mutualist *S. symbiotica* strain in *Acyrthosiphon pisum* can protect the host against parasitoid wasps (Oliver et al. 2006). In some aphid species, *S. symbiotica* is also believed to be assuming the role of a co-obligate symbiont to compensate for declining efficiency of *Buchnera* as the ancestral primary nutritional symbiont (Monnin et al. 2020).

In this study, we identify and genomically characterise a new *Serratia* taxon associated with populations of the redlegged earth mite, *Halotydeus destructor* (Tucker, 1925) (Arachnida: Trombidiformes: Penthaleidae) in its invasive Australian range. *Halotydeus destructor* is a major invasive agricultural pest of grain and pasture systems in Australia that causes major yield losses (Ridsdill-Smith 1997). The detection of *Serratia* reads in previously collected population genomic data of *H. destructor* provided an opportunity to investigate a bacterial association in this mite, which has had no formal characterisation of its bacterial symbionts to date. Notably, in other mites and ticks, all recorded *Serratia* species are free-living, which include *S. marcescens*, *S. liquefaciens*, and *S. odorifera* (Chaisiri et al. 2015; Hubert et al. 2016; Castro-Saines et al. 2022; Sumner-Kalkun et al. 2023).

Here, we use metagenomic approaches to assemble the genome of the *H. destructor-*associated *Serratia* species and test for the genomic signatures of host adaptation. We predicted that host adaptation in our new *Serratia* taxon would be evident from a reduction in genome size and coding capacity, an increased abundance of mobile genetic elements, greater genomic rearrangement, changes in overall gene content, and remodelling of the repertoire of virulence factors, relative to free-living *Serratia*. We also reanalysed previously generated sequencing data to map the occurrence of our new *Serratia* species across *H. destructor* populations to infer its broader ecological relevance in the invasive range of its host.

## Results

### Two closely related *Serratia* genomes assembled from *H. destructor*

We assembled two *Serratia* genomes from *H. destructor*. These genomes are broadly similar, with modest differences in genome size and predicted coding capacity (Table 1). Each final assembly was approximately 3.7 Mb in size, comprised 22 contigs, and had an N50 ranging from 246 kb to 248 kb. The GC content of both genomes was approximately 53%, and 3,514 and 3,541 coding sequences were predicted for the two strains, respectively. Recovery of 98.4% of single-copy bacterial BUSCOs indicated that these *Serratia* assemblies were nearly complete (Table 1). The near identical assembly statistics and high completeness of the two genomes provided a consistent basis for evaluating lineage-level genomic features.

**Table 1.**
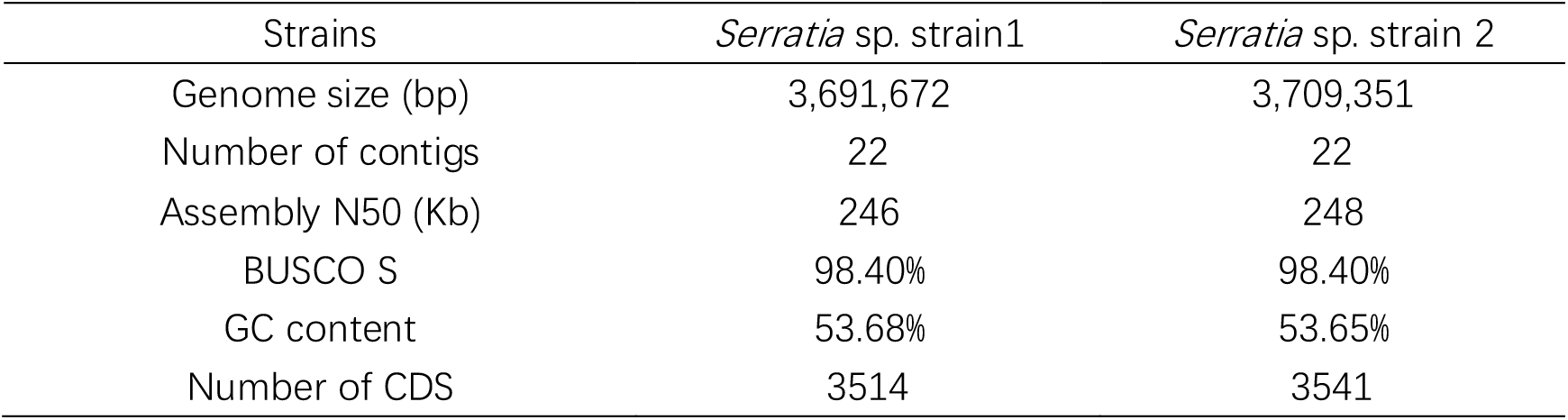
Genomes of *Serratia* sp. strains from *H. destructor*.

| Strains | *Serratia* sp. strain1 | *Serratia* sp. strain 2 |
|---|---|---|
| Genome size (bp) | 3,691,672 | 3,709,351 |
| Number of contigs | 22 | 22 |
| Assembly N50 (Kb) | 246 | 248 |
| BUSCO S | 98.40% | 98.40% |
| GC content | 53.68% | 53.65% |
| Number of CDS | 3514 | 3541 |

### *H. destructor*-associated *Serratia* is likely a unique taxon with genomic signatures of host adaptation

Our phylogeny of *Serratia* comprised the two *H. destructor*-associated *Serratia*, 13 *Serratia symbiotica* strains, and 24 free-living *Serratia* species (Fig. 1A). Two major lineages emerged: all *S. symbiotica* strains formed a distinct clade separated from the free-living species and were relatively distant from their closest free-living relatives.

**Fig. 1.**
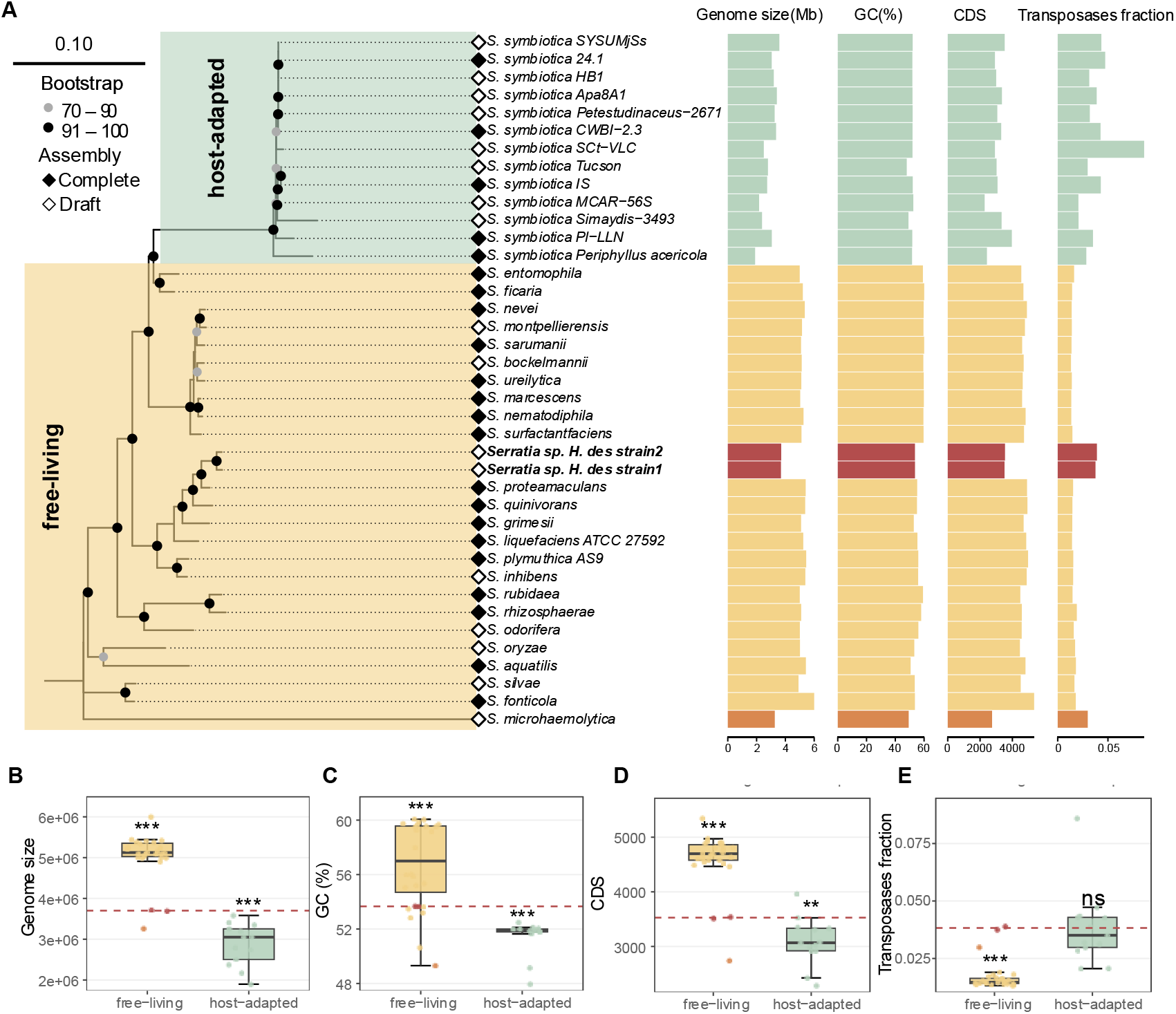
Phylogenetic placement and genomic characteristics of the *H. destructor*-associated *Serratia* sp. (A) Maximum-likelihood phylogeny inferred from an alignment of 658 core genes from 39 *Serratia* strains using 1,000 ultrafast bootstrap replicates. Free-living *Serratia* are shown in yellow, host-adapted *S. symbiotica* strains in green, the two *H. destructor*-associated *Serratia* sp. strains are in red, and the basal *Serratia* taxon in the displayed tree in orange. Black and grey circles indicate bootstrap support values of 91-100% and 70-90%, respectively. Filled and open diamonds indicate complete and draft genome assemblies, respectively. Horizontal bar plots show genome size in megabases, GC content, number of predicted coding sequences (CDS), and transposase fraction for each genome. The transposase fraction was calculated as the number of genes annotated as transposases divided by the total number of CDS. (B-E) Boxplots summarising distributions of genome size (B), GC content (C), CDS number (D), and transposase fraction (E) between free-living and host-adapted *Serratia*. Individual points represent genomes, and the red dashed line indicates the mean value of the two *H. destructor*-associated *Serratia*. Two-sided one-sample Student’s *t*-tests were used to determine whether the mean of each free-living or host-adapted group differed from the corresponding mean value of *Serratia* sp. Significance levels are denoted as follows: ns, not significant (P ≥ 0.05); *, P < 0.05; **, P < 0.01; and ***, P < 0.001) *Serratia* sp.

Our two *H. destructor*-associated *Serratia* genomes were placed within the free-living group, clustering near the *S. liquefaciens* complex. Measures of whole-genome average nucleotide identity (ANI) criteria revealed that both *H. destructor*-associated *Serratia* strains had less than 95% identity with all described *Serratia* species included in our analysis (Table S1). These findings suggest that our *H. destructor*-associated *Serratia* is a unique taxon and possibly a new species.

Genomic characteristics suggest that *H. destructor*-associated *Serratia* sp. exhibited quite different patterns from free-living species and has similarities to host-adapted *S. symbiotica*. Free-living *Serratia* tended to have genomes that were larger, had a higher GC content, encoded more genes, and had fewer transposases, relative to the host-adapted *S. symbiotica* (Fig. 1 B−E). We found that the *H. destructor*-associated *Serratia* sp. showed genomic characteristics intermediate between free-living and host-adapted lineages, with significant differences from both groups for every characteristic except transposase fraction, which did not differ significantly from host-adapted *Serratia* (Fig. 1 B−E). Surprisingly, we observed similar genomic changes in the genome of *S*. *microhaemolytica*, which is the basal lineage of our tree.

Structural comparisons using synteny analysis also suggest that our *H. destructor*-associated *Serratia* sp. exhibits less synteny relative to other free-living *Serratia* (Fig. 2). For reference, we compared the free-living *S. proteamaculans* and *S. ficaria*, which are quite distantly related: *S. proteamaculans* is closely related to our *H. destructor Serratia* sp., and *S. ficaria* is more closely related to *S. symbiotica*. We observed greater collinearity between *S. proteamaculans* and *S. ficaria* than between our *H. destructor Serratia* sp. and *S. proteamaculans*, which implies genomic rearrangements have occurred in the *H. destructor*-associated *Serratia* sp.

**Fig. 2.**
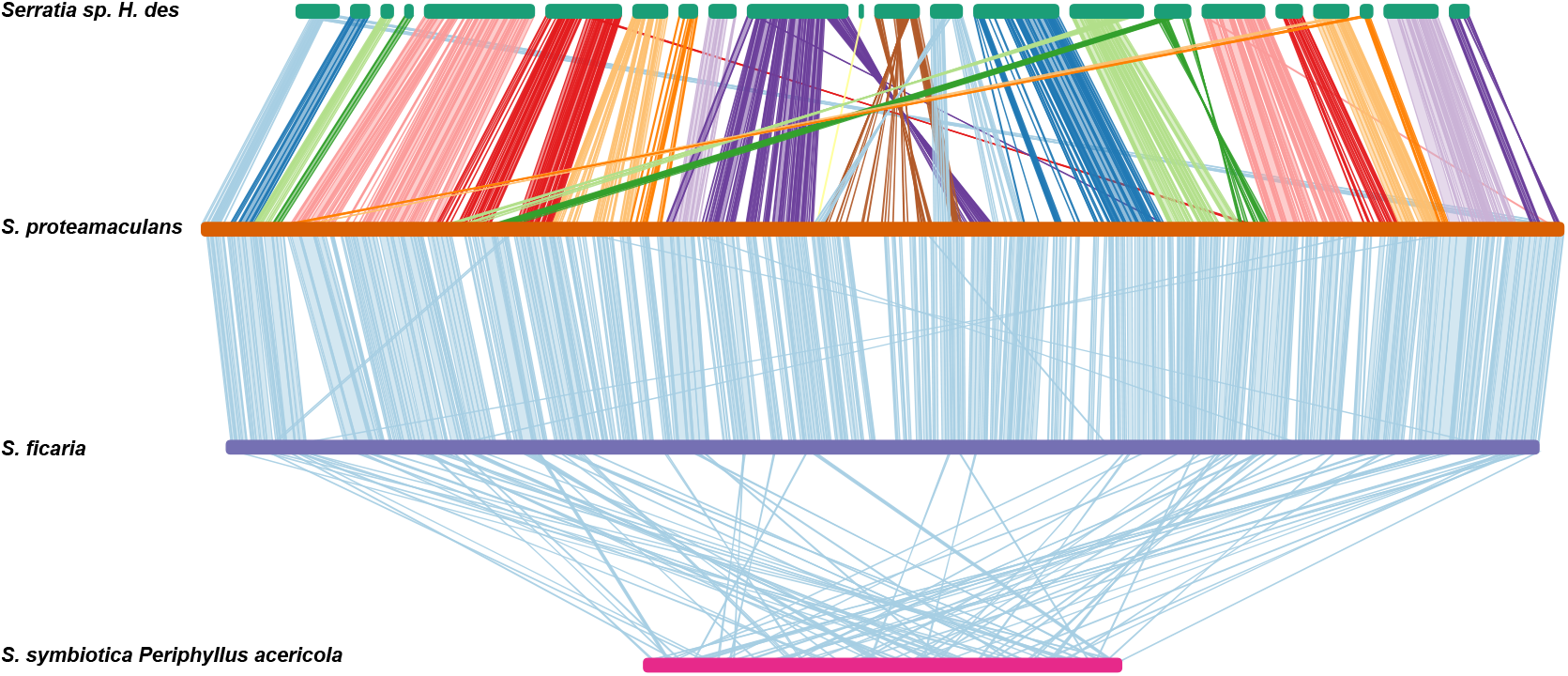
Whole genome synteny comparison among representative *Serratia* strains. Four representative strains were selected to represent phylogenetic and lifestyle contrasts: the *H. destructor*-associated *Serratia* sp. and its close free-living strain *S. proteamaculans*, as well as *S. symbiotica Periphyllus acericola* and its close free-living strain *S. ficaria*. The contigs of *Serratia* sp. were reordered based on the *S. proteamaculans* genome as a reference. Connecting ribbons represent homologous genomic regions of at least 1 kb; crossing or displaced ribbons indicate inversions and other chromosomal rearrangements.

Collectively, our results suggest *H. destructor*-associated *Serratia* sp. is an undescribed species. Although this lineage retains phylogenetic similarity to free-living *Serratia*, it shows genomic signatures typical of host adaptation. These data might thus suggest a possible independent transition toward a host-adapted symbiosis in the main free-living *Serratia* clade.

### Gene content in our *H. destructor Serratia* sp. is distinct from other *Serratia*

Orthologous gene content was used to further describe differences between *H. destructor*-associated *Serratia* sp. and other *Serratia*. We first compared broad functional profiles based on four major Clusters of Orthologous Genes (COG) categories. Generally, free-living and host-adapted *Serratia* groups showed contrasting gene-content distributions across several of these categories. However, *S. microhaemolytica* appears to be an exception, as its gene proportions across all four categories differ from those of other free-living *Serratia*. Relative to the host-adapted group, free-living *Serratia* genomes tended to contain a lower proportion of genes assigned to cellular processes and signaling (Fig. 3A), but a higher proportion of genes assigned to metabolism and poorly characterized functions (Fig. 3C, D). By contrast, genes assigned to information storage and processing showed no clear separation between the free-living and host-adapted groups (Fig. 3B).

**Fig. 3.**
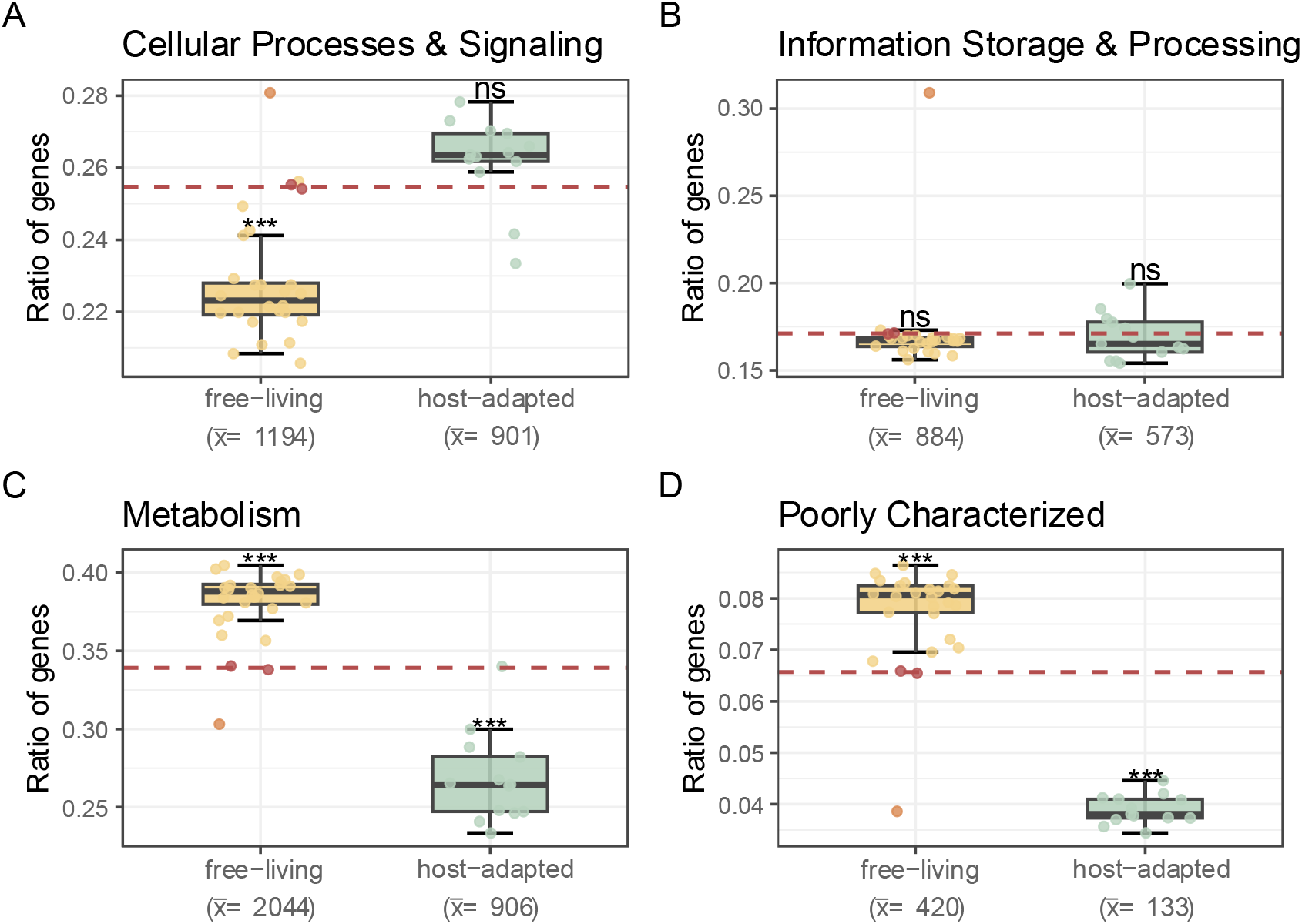
Broad COG functional profiles of free-living, host-adapted, and *H. destructor*-associated *Serratia* sp. (A–D) Proportions of genes assigned to the four broad COG functional classes: cellular processes and signalling (A), information storage and processing (B), metabolism (C), and poorly characterized functions (D). For each genome, the proportion was calculated as the number of genes assigned to the corresponding class divided by the total number of genes. Mean gene counts per genome for each group are shown in parentheses below the corresponding labels. Yellow and green points represent individual free-living and host-adapted *Serratia* genomes, respectively; red points represent the two *Serratia* sp. strains, and the red dashed line indicates their mean. Two-sided one-sample Student’s *t*-tests were used to determine whether the mean of each free-living or host-adapted group differed from the corresponding mean value of *Serratia* sp. Significance levels are denoted as follows: ns, not significant (*P* ≥ 0.05); *, *P* < 0.05; **, *P* < 0.01; and ***, *P* < 0.001)

Across these categories, our *H. destructor*-associated *Serratia* sp. generally occupied positions between the two comparison groups. The proportion of cellular processes and signaling genes in *H. destructor*-associated *Serratia* sp. exceeded that observed among free-living *Serratia* (*t* = −8.064, df = 23, P < 0.001), whereas it did not differ significantly from that of host-adapted *Serratia* (Fig. 3A). The proportion of genes assigned to information storage and processing did not differ significantly between *H. destructor Serratia* sp. and either group (Fig. 3B). The proportion of metabolism genes in *H. destructor*-associated *Serratia* sp. was significantly lower than free-living *Serratia* (*t* = 10.191, df = 23, P < 0.001) and significantly larger than host-adapted *Serratia* (*t* = −8.811, df = 12, P < 0.001) (Fig. 3C). Finally, the proportion of poorly characterized genes in *H. destructor*-associated *Serratia* sp. was significantly lower than free-living *Serratia* (*t* = 6.077, df = 23, P < 0.001) and significantly higher than host-adapted *Serratia* (*t* = −33.809, df = 12, P < 0.001) (Fig. 3D).

We next identified the conserved genes across both the free-living and host-adapted *S. symbiotica*, as well as genes conserved within each group, and then compared the presence-absence profiles of our *H. destructor*-associated *Serratia* sp. against these sets (Fig. 4). Our *H. destructor*-associated *Serratia* sp. shares genes that are conserved across *Serratia* and conserved within the free-living lineage. However, there did appear to be a general loss of the conserved free-living lineage genes. Although our *Serratia* sp. does not generally share genes in the conserved *S. symbiotica* cluster, it does have its own unique gene cluster. This cluster corresponds mainly to mobile elements, such as prophage and transposon related genes, and genes related to intracellular trafficking, secretion, and vesicular transport (Fig. S1). The unique mobile elements are consistent with the elevated transposase fraction observed in the genome and may contribute to genome rearrangements. The enrichment of functions in intracellular trafficking, secretion, and vesicular transport might suggest expansion of functions associated with host interaction. If *H. destructor*-associated *Serratia* sp. is shifting to a host-adapted lifestyle, these gene patterns might suggest that they retain some core free-living genes whilst evolving a unique independent set of genes different to those evolved in *S. symbiotica*.

**Fig. 4.**
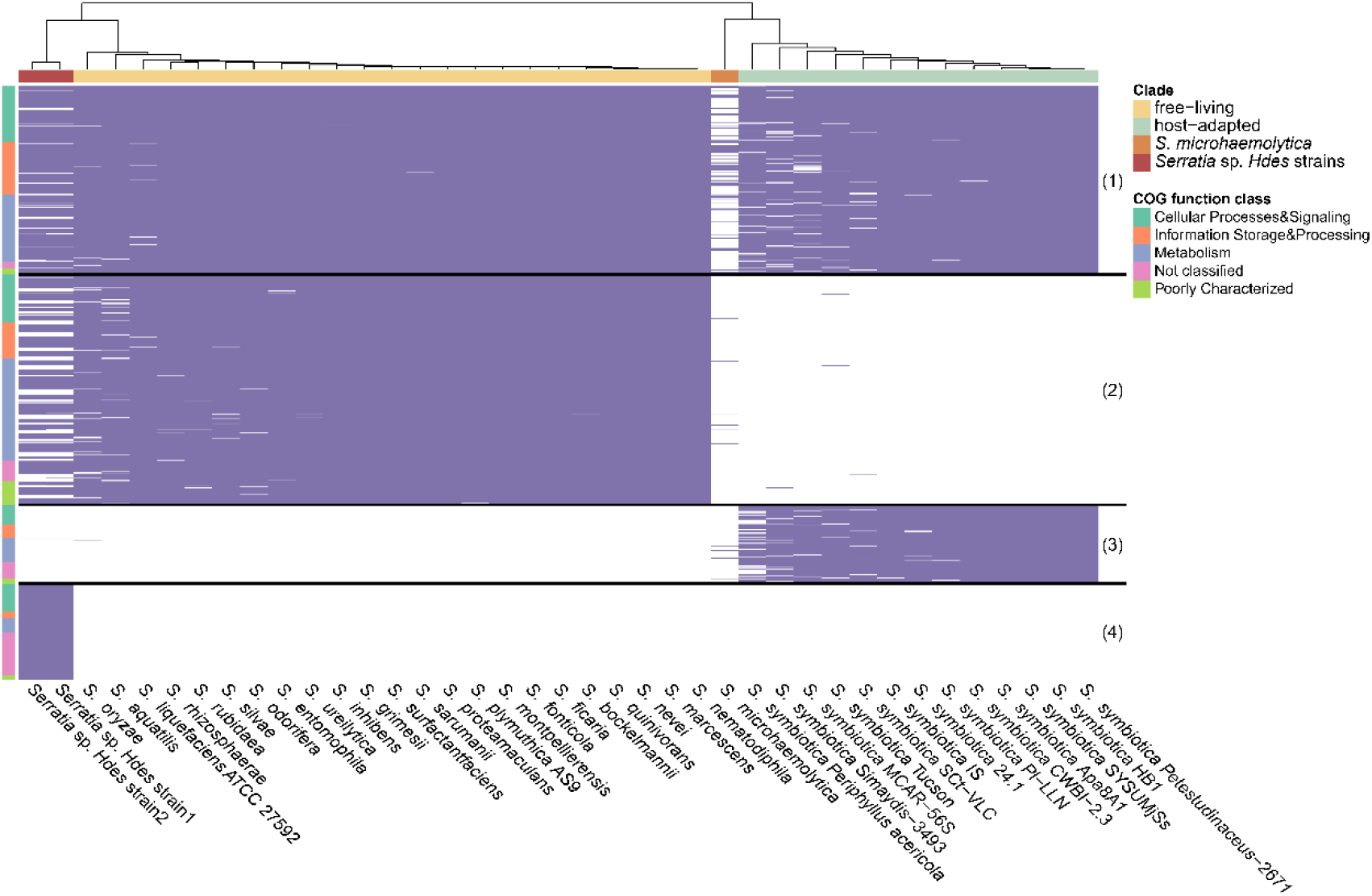
Orthologous genes presence and absence across *H. destructor*-associated *Serratia* sp., free-living *Serratia* and host-adapted *Serratia* in COG categories. Columns represent individual genomes and rows represent orthologous gene families selected, horizontal lines separate sets of key genes related to (1) conserved genes across both free-living and host-adapted groups, (2) the free-living and (3) the host-adapted group-specific conserved genes, as well as (4) the unique genes only present in *H. destructor*-associated *Serratia* sp.. Purple cells indicate gene presence, and white cells indicate gene absence. The dendrogram above shows clustering of strains based on the gene’s presence and absence pattern. The top bar identifies free-living taxa in yellow, host-adapted taxa in green, the two *Serratia* sp. strains in red, and the basal *Serratia* taxon in orange. The annotation bar on the left indicates the broad COG functional class assigned to each gene family; families without a COG assignment are designated as not classified.

### Remodelling of virulence factors supports a lifestyle shift in *H. destructor-associated Serratia* sp

We further compared virulence factor (VF) repertoires across all *Serratia* genomes to identify changes in gene profiles that might be associated with host adaptation (Fig. 5). VF genes with considerable decreases in *H. destructor*-associated *Serratia* sp. included those involved in adherence and motility, whereas effector delivery system genes increased. This pattern is consistent with the loss of genes associated with free-living functions and the enrichment of genes involved in host interactions. The overall gene proportion of VF repertoires in *H. destructor*-associated *Serratia* sp. showed that the two strains did not resemble the conserved VF profile characteristic of most free-living *Serratia* or host-adapted *S. symbiotica.* Instead, their VF profiles were largely intermediate, as visualised in a principal component analysis (Fig. 6A). Measures of VF profile distance within each clade indicated that free-living *Serratia* had, on average, more conserved gene contents than *S. symbiotica* (Fig. 6B) (PERMDISP analysis, F = 12.48, P = 0.001), but *H. destructor*-associated *Serratia* sp. was more divergent within the free-living clade.

**Fig. 5.**
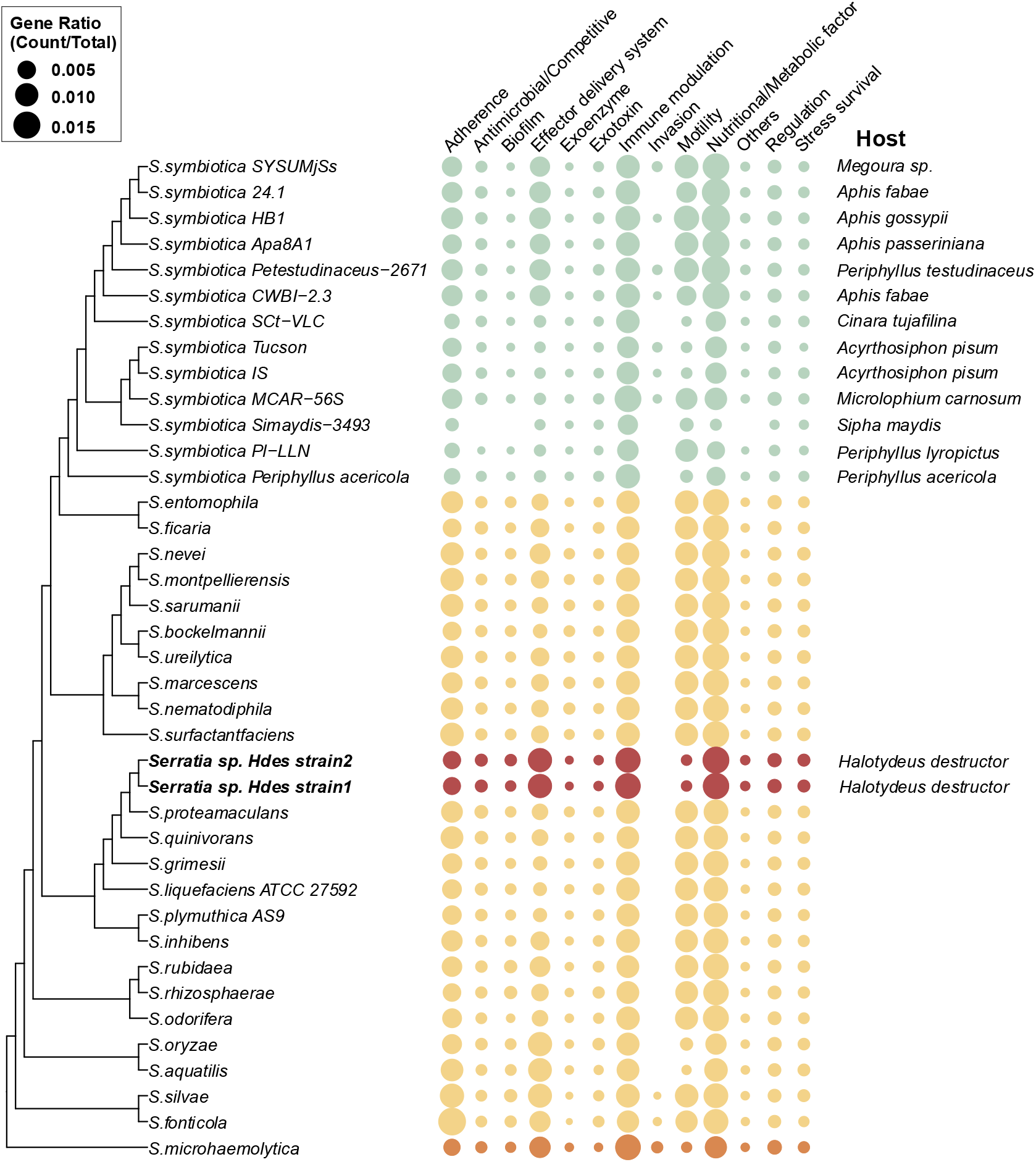
Remodelling of virulence factor repertoires across free-living and host-adapted *Serratia* lineages. Bubble sizes indicate the proportion of genes assigned to each virulence factor (VF) functional category relative to the total number of predicted coding sequences in each genome. Strains are ordered according to the core-genes phylogeny. Free-living *Serratia* are highlighted in yellow, host-adapted *Serratia* in green, *H. destructor*-associated *Serratia* sp. in red. and the basal *Serratia* taxon in the phylogenetic tree in orange. Functional categories were assigned according to VFDB annotations. Host species of host-adapted lineages are listed on the right.

**Fig. 6.**
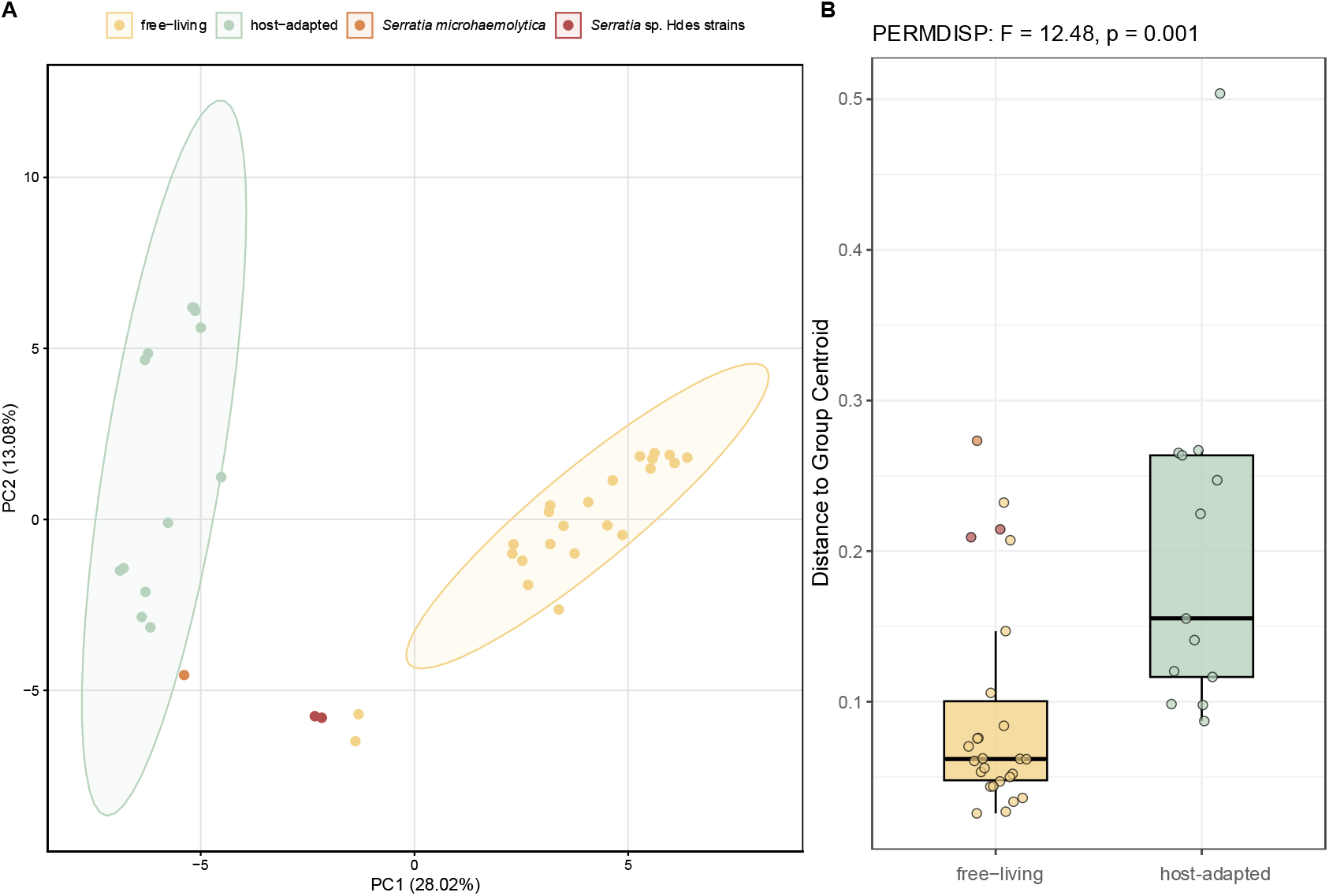
Functional divergence of virulence factors in *Serratia* species. (A) Principal component analysis (PCA) based on the presence and absence of virulence factors matrix. (B) Within-group dispersion of VF repertoires estimated using Bray-Curtis distances (PERMDISP). Statistical parameters (F-value and associated p-value) were derived from a permutation test with 999 permutations.

We also note that the remodelling of virulence factors occurred in the genome of *S*. *microhaemolytica*.

### *Serratia* sp. is broadly distributed across *H. destructor* populations in Australia

We used our *Serratia* sp. genomes to genomically screen for infections across 190 populations sampled across Australia between 2018 to 2023. Using a combination of read depth and genomic coverage, we could confidently identify infections across 72 populations, distributed throughout Australia (Fig. 7). The repeated detection and broad geographic distribution of *Serratia* sp. across *H. destructor* populations are consistent with a general association between the bacterium and its host across its invasive Australian range.

**Fig. 7.**
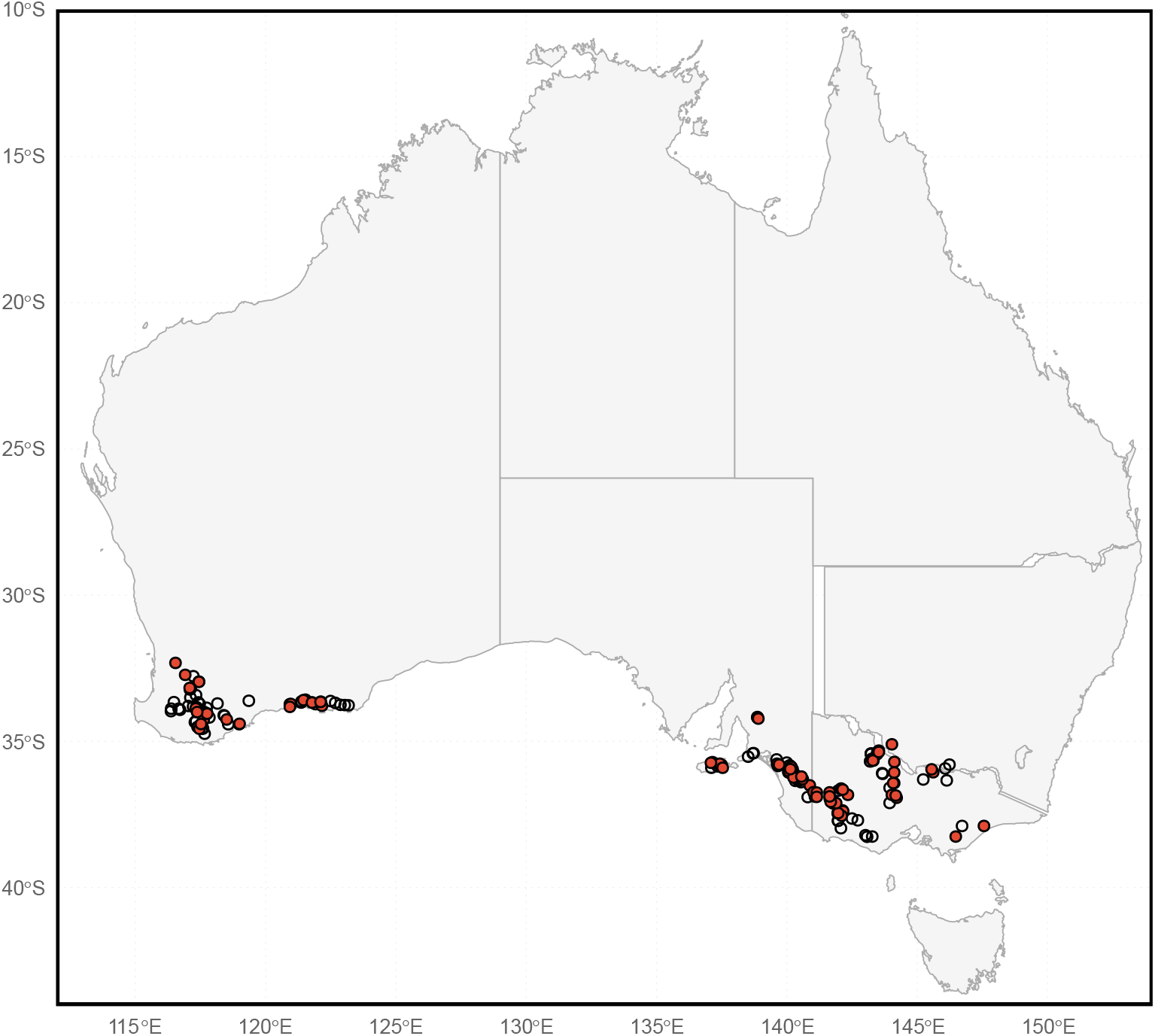
Geographic distribution of *Halotydeus destructor* populations with genomic evidence of infection by *Serratia* sp. across Australia. Pooled whole genome sequencing data from 190 *H. destructor* populations sampled between 2018 and 2023 were screened by mapping non-host reads to the reconstructed *Serratia* sp. genome. Red circles indicate the 72 populations classified as infected; populations below this threshold are open circles.

## Discussion

Our newly identified *Serratia* sp. from the trombidiform mite, *H. destructor*, represents a potentially parallel transition to a host-adapted lifestyle outside the *S. symbiotica* lineage. Our phylogenetic reconstruction of *Serratia* species is consistent with previous work (Williams et al. 2022), and *H. destructor*-associated *Serratia* sp. sits within the free-living lineages apart from the phylogenetically distant *S. symbiotica*. Nevertheless, our *Serratia* sp. exhibits multiple genomic signatures typically associated with adaptation to arthropod hosts, including genome reduction, elevated transposase fraction, chromosomal rearrangements, shifts in gene proportion across COG categories, and remodelling of virulence factor repertoires. Similar genomic changes have been documented in *S. symbiotica* (Burke and Moran 2011; Manzano-Marín et al. 2012; Renoz et al. 2021) and across other bacterial symbioses (Akman et al. 2002; Degnan et al. 2010; Otero-Bravo et al. 2018). However, even compared to the *S. symbiotica* strains with less genome reduction, which are thought to represent early stages of host adaptation (Sabri et al. 2011; Renoz et al. 2021), the larger genome, higher GC content, and broader coding capacity of *H. destructor*-associated *Serratia* sp. suggests it may be in an earlier stage of transition along the continuum from free-living to host-adapted genomes.

Our new *H. destructor*-associated *Serratia* sp. represents the first putatively host-adapted *Serratia* taxon characterised in Acari (mites and ticks). Within Trombidiformes, *Serratia* was previously detected in the eggs of *Tetranychus urticae*, but with much lower abundance in adults; thus the authors suggested that its presence was likely resulting from environmental contamination (Merlin et al. 2023). In other mite orders, *S. marcescens* has been reported in the parasitic mite *Psoroptes ovis* (Hogg and Lehane 1999), *S. marcescens* and *S. liquefaciens* have been isolated from the Sarcoptiformes mite *Tyrophagus putrescentiae* (Smrž et al. 2016), and *S. marcescens* and *S. odorifera* have been identified in predatory mites (Mesostigmata) (Sumner-Kalkun et al. 2023). In ticks (Ixodida), *S. marcescens* has also been identified in the cattle tick (Andreotti et al. 2011). However, these studies have primarily documented bacterial occurrence rather than providing evidence associated with adaptation to the host. This is unsurprising because host-adapted *Serratia* strains have been assumed to be restricted to *S. symbiotica*. Although *S. symbiotica* has occasionally been detected outside aphids, such as in aphid-tending ants (Renoz et al. 2019), it has not been described in mites (or ticks).

The genomic changes found in our study may reflect a broader adaptation process of ecological specialization. Changes can arise as bacteria transition from diverse environments to more restricted ecological niches, whether within a host or in specialized free-living environments. This process could be affecting the genome of the free-living *Serratia microhaemolytica*, which despite not adapting to a host nevertheless shows genome reduction, transposon expansion, shifts in functional gene composition, and virulence factor remodeling (Figs. 1B-E, 3-6). This species was isolated from an artificial lake and proved to be culturable, with no evidence of a specific host association (Wang et al. 2019). Its genomic changes may therefore reflect ecological specialization in a free-living environment rather than host adaptation. Genome reduction associated with ecological specialization has been documented in other free-living bacteria such as *Prochlorococcus* and *Idiomarina* (Dufresne et al. 2005; Qin et al. 2019). By contrast, we have found a high prevalence of *H. destructor Serratia* sp. Across host populations from a variety of locations, and years, supporting the notion that the new *Serratia* species represents a tight host association rather than being associated with other environmental factors.

Recovering genomes from metagenomic data containing two coexisting strains is challenging. Here we showed that cross sample abundance variation can serve as an additional signal to resolve strain-specific assemblies. Although our assemblies may underestimate some strain-specific structures and contain minor Nanopore-derived base errors, they retain the major genomic features needed for comparative analyses. As genome reduction in *H. destructor*-associated *Serratia* sp. is less extensive than in culturable *S. symbiotica*, future work should assess its ability to be cultured, as clonal isolates would enable complete, high-quality genome assembly.

Although genomic approaches provide valuable insights into transitions towards host-adapted lifestyles, experimental evidence is crucial for understanding how symbiotic bacteria influence the evolutionary ecology of their hosts. *Halotydeus destructor* is a widespread invasive agricultural pest in Australia, where pesticide resistance has become an increasing management concern (Thia et al. 2023). *Serratia* could have contrasting effects on the hosts’ pesticide resistance. *Serratia symbiotica* can increase susceptibility to several insecticides in the pea aphid *Acyrthosiphon pisum* (Skaljac et al. 2018), while several free-living *Serratia* species are involved in the degradation or detoxification of pesticides (Xia et al. 2023; Cycoń et al. 2009; Alatassi et al. 2025), and may therefore increase host pesticide resistance. With our genomic evidence suggesting *H. destructor*-associated *Serratia* sp. occupies an intermediate evolutionary position between free-living *Serratia* and *Serratia symbiotica*, it would be interesting to see how it affects host pesticide resistance.

## Conclusion

Our study highlights the value of metagenomic and genomic approaches for characterizing bacterial symbionts and their potential roles in the evolutionary continuum from free-living to a host-adapted lifestyle. Broadly, our *H. destructor*-associated *Serratia* sp. shows that host adaptation in the genus *Serratia* is not confined to *Serratia symbiotica*, and the new species identified here may provide an intermediate stage in symbiosis evolution. Despite its widespread occurrence across *H. destructor* populations, the effects of this new species on host biology remain unknown. A critical next step will be to derive experimental data to link putative phenotypic and fitness effects of the *H. destructor*-associated *Serratia* sp. to its mite host.

## Methods

### Genomic sequence data

All genomic sequence data was generated in a previous study (Thia et al. 2025). The long reads of *H. destructor* population RLEM102 were used to assemble draft *Serratia* genomes. The short reads of population RLEM102 were used for metagenome binning. The short reads of population RLEM130 were used to separate the two assemblies, as described in the following section.

### Metagenome assembly and annotation

*Serratia* metagenomic assembly was performed using Flye v2.9.2 with the “--meta” mode (Kolmogorov et al. 2020). High redundancy in this preliminary assembly suggested the presence of more than one closely related *Serratia* strain. We determined that these were likely two different strains of the same taxon, rather than different taxa, based on analysis of the 16S rRNA sequence (Fig S2). Two putative genomes of *H. destructor*-associated *Serratia* were obtained by phasing the long-reads assembly. The preliminary assembly was first binned using metaWRAP v1.3, after which redundant contigs were manually removed to curate the assembly. Long reads were aligned back to the binned assembly using minimap2 v2.26 (Li 2018). Variants were called using freebayes v1.3.6 (Garrison and Marth 2012), filtered using BCFtools v1.15.1 (Danecek et al. 2021), and phased using HAPCUT2 v1.3.4 (Edge et al. 2017). Haplotype-resolved assemblies were then extracted from the phased VCF using BCFtools v1.15.1.

To further assign phased contigs to strain-specific assemblies, we used differences in relative depth across additional *H. destructor* populations. Because the relative abundance of the two *Serratia* strains varies among populations, contigs originating from the same strain are expected to have similar depth, whereas corresponding contigs from the two strains should differ in depth. We therefore mapped reads from 10 randomly selected populations and compared their sequencing depths across populations. To minimize cross-mapping and ensure the accuracy of haplotype-specific depth estimation, we applied more stringent read mapping using Bowtie (Langmead et al. 2009) with the parameter “-v 1”. Contig depths were estimated using the jgi_summarize_bam_contig_depths from MetaBAT v2.15, which filters out unmapped or low-quality reads and calculates the mean depth over the entire length of the contig using only reliably mapped reads (Kang et al. 2019). For the subsequent contig assignment, we selected a specific population (RLEM130) that exhibited a substantial depth difference between the two haplotype genomes (Table S2). The two strain-specific assemblies were obtained by reassigning the contigs with similar depth to the same assembly.

Completeness of the strain-specific assemblies was assessed with BUSCO v5.4.5 (Manni et al. 2021). Annotations of the genomes were performed using Prokka v1.14.5 (Seemann 2014).

### Phylogenomic analysis, genomic characteristics, and genomic synteny

Phylogenomic analysis was used to understand the taxonomic context of our *H. destructor*-associated *Serratia*. We compiled a set of *Serratia* genomes that were downloaded from the NCBI RefSeq database (Table S3). In this dataset, we included one representative genome from each available free-living *Serratia* species to capture broad taxonomic diversity, together with *S. symbiotica* strains representing different stages of host adaptation. For this comparison, *Serratia* species capable of a free-living lifestyle were classified as free-living regardless of reported host associations, whereas *S. symbiotica* strains were classified as host-adapted. The plasmid sequences of genomes were removed prior to analysis. All genomes were re-annotated using Prokka v1.14.5 to ensure annotation consistency across datasets. Orthologous genes among *Serratia* genomes were identified using Panaroo v1.5.2 (Tonkin-Hill et al. 2020) in moderate cleaning mode. Initial gene clustering used a sequence identity threshold of 95% and a length similarity cutoff of 0.95, restricting clustering to sequences with no more than approximately 5% difference in length. Gene-family clustering was performed using a protein sequence identity threshold of 50%. Core genes were defined as those present in at least 95% of genomes. Maximum-likelihood phylogenetic trees were inferred from core-gene alignments using IQ-TREE v2.1.2 with 1,000 ultrafast bootstrap replicates (Minh et al. 2020). The tree was rooted using *Yersinia enterocolitica* as an outgroup (Fig S3), as it is closely related to the genus *Serratia*.

To test the hypothesis that the *H. destructor*-associated *Serratia* represents a distinct taxon, pairwise average nucleotide identity (ANI) was calculated from whole-genome sequences using FastANI v1.3.4 (Jain et al. 2018). We applied the commonly used ANI species boundary of approximately 95% to determine whether strains belonged to the same species or represented distinct taxa (Richter and Rosselló-Móra 2009).

Next, we examined the genomic characteristics across all *Serratia* species to infer the potential lifestyle of the *H. destructor*-associated *Serratia*. Relative to free-living species, bacteria that have evolved more intimate symbioses with their host are expected to have a smaller genome and less coding capacity, lower GC content, enrichment of mobile elements and chromosomal rearrangements. We summarised these characteristics separately for the previously sequenced free-living and host-adapted *Serratia*, respectively, and tested for significant differences against our *H. destructor*-associated *Serratia* using *t*-tests.

Finally, we examined patterns of genomic synteny among a select number of genomes to further evaluate evidence of a lifestyle shift. We measured the genomic synteny between *H. destructor*-associated *Serratia* and its closely related free-living species, *S. proteamaculans*, and to a free-living species that was more closely related to the host-adapted species, *S. ficaria*, and to the host-adapted *S. symbiotica Periphyllus acericola*. Genomic sequences were aligned using MUMmer v4.0.0 (Marçais et al. 2018) and alignments shorter than 1 kb were excluded. Genome synteny was visualized using NGenomeSyn v1.4.3 (He et al. 2023).

### Gene content analyses

Patterns of gene content were used to understand the evolutionary uniqueness of our *H. destructor*-associated *Serratia* and to further assess its shift toward host adaptation. First, we broadly assessed differences in functional composition among *Serratia* taxa using the Clusters of Orthologous Genes (COG) approach. Orthologous genes from Panaroo were aligned against the COG database (Galperin et al. 2025) using DIAMOND v2.1.0 (Buchfink et al. 2015) with an E-value threshold of 1e−5. COG categories were then used to create a gene presence and absence matrix. Genes assigned to multiple COG categories were counted separately for each category in later statistical analyses, following the approach used by another study (Cohen et al. 2011). These annotations were used to summarize the proportional representation of genes in COG functional categories. To avoid bias caused by differences in genome size, the number of genes in each category was normalized to the total number of genes in each genome. We tested the hypotheses that our *H. destructor*-associated *Serratia* had significantly different functional compositions to free-living and host-adapted *Serratia* using a *t*-test, as implemented in R.

Next, we identified functional sets of genes that were shared and unique to free-living and host-adapted *Serratia* and used these as a reference to compare with our *H. destructor*-associated *Serratia*. Because the groups contained only a modest number of genomes, we used a predefined absolute tolerance of two genomes as an operational criterion for identifying near-ubiquitous gene families, rather than assigning them to conventional percentage-based core-genome categories. Based on this, we listed (1) conserved genes across both free-living and host-adapted groups when present in all genomes while allowing up to two absences, as well as (2) the free-living and (3) the host-adapted group-specific conserved genes, as genes absent from fewer than two strains within the focal group and present in fewer than two strains of the comparison group. *H. destructor*-associated *Serratia* were then independently projected onto these predefined gene sets, allowing a direct comparison. Finally, we listed (4) the unique genes present only in *H. destructor*-associated *Serratia* genomes.

Finally, we examined the composition of virulence factors across species to assess the shift of *H. destructor*-associated *Serratia* toward a host-adapted lifestyle. Protein sequences of *Serratia* strains were queried against the VFDB protein database (Zhou et al. 2025) using DIAMOND v2.1.0 with an E-value threshold of 1e−5, a minimum identity threshold of 50%, a minimum alignment coverage threshold of 80%, and retention of the best hit only. Then a presence and absence matrix of virulence factors across all *Serratia* was constructed for further analysis. Composition of virulence factors was visualised by aligning genes against the core gene phylogeny and by performing a PCA on the presence-absence matrix in R. Distances in composition within the free-living and host-adapted groups were calculated using Bray-Curtis distances and tested using PERMDISP with 999 permutations. As in the COG analyses, we normalized the counts of virulence factors by the total gene count in each genome.

### Inferring infection status within and among populations

To characterise the spatial and temporal distribution of our *Serratia* taxon across populations of *H. destructor*, we re-analysed short-read pool-seq data from 190 populations sampled across years and locations in Australia. These pool-seq data came from (Thia et al. 2025). We mapped short-read sequences against the *H. destructor* genome using Bowtie2 v2.4.5 (--very-sensitive) and extracted those reads that did not align. These were then mapped against our new *H. destructor*-associated *Serratia* assemblies to obtain estimates of read depth and genomic coverage using a custom Perl script, with a minimum of two reads required to support a site. We inferred strong evidence for infection by examining the relationship between genomic coverage and normalized read depth (Fig S4).

## Data availability

The draft genome assemblies of *Serratia* sp. have been submitted to NCBI GenBank and are currently being processed; the corresponding accession numbers will be provided upon assignment (private review link: https://figshare.com/s/6f57f8173b541f9746e9). The long reads and short reads data used in this study are deposited under the NCBI BioProject PRJNA756307 and presented in Table S4.

## Acknowledgments

Genomic and molecular work was supported through an investment from the Grains Research and Development Corporation (UOM1906-002RTX; UOM2404-006RT) and Hort Innovation (ST23002) as part of the ‘Australian Grain and Horticulture Pest Innovation Program’ to AAH, with additional investment from the University of Melbourne and Cesar Australia. This research was supported by The University of Melbourne’s Research Computing Services and the Petascale Campus Initiative. TLS was supported by an ARC DECRA (DE230100257).

## Author contributions

JC, AAH, TLS and JAT conceived and designed the study. JC performed the bioinformatic analyses. JAT provided the short-read sequencing data and contributed to short-read alignment and processing. SBS and NDY contributed to the generation of Nanopore long-read data. TLS and AAH acquired funding to support JC. JC drafted the manuscript. AAH, TLS and JAT reviewed and edited the manuscript.

## AI use statement

ChatGPT (OpenAI) was used to check English spelling and grammar, and to assist with implementing and refining R scripts based on analytical workflows and statistical approaches designed by the author.

